# Beyond microtubules: The cellular environment at the endoplasmic reticulum attracts proteins to the nucleus, enabling nuclear transport

**DOI:** 10.1101/2024.01.12.575351

**Authors:** Seok Joo Chae, Dae Wook Kim, Oleg A. Igoshin, Seunggyu Lee, Jae Kyoung Kim

**Affiliations:** Department of Mathematical Sciences, KAIST, Daejeon 34141, Republic of Korea; Biomedical Mathematics Group, Pioneer Research Center for Mathematical and Computational Sciences, Institute for Basic Science, Daejeon 34126, Republic of Korea; Department of Mathematics, University of Michigan, Ann Arbor, MI 48109, USA; Department of Bioengineering, Rice University, Houston, TX 77005, USA; Center for Theoretical Biological Physics, Rice University, Houston, TX 77005, USA; Department of Chemistry, Rice University, Houston, TX 77005, USA; Department of Biosciences, Rice University, Houston, TX 77005, USA; Division of Applied Mathematical Sciences, Korea University, Sejong 30019, Republic of Korea

## Abstract

All proteins are translated in the cytoplasm, yet many, including transcription factors, play vital roles in the nucleus. While previous research has concentrated on molecular motors for the transport of these proteins to the nucleus, recent observations reveal perinuclear accumulation even in the absence of an energy source, hinting at alternative mechanisms. Here, we propose that structural properties of the cellular environment, specifically the endoplasmic reticulum (ER), can promote molecular transport to the perinucleus without requiring additional energy expenditure. Specifically, physical interaction between proteins and the ER impedes their diffusion and leads to their accumulation near the nucleus. This result explains why larger proteins, more frequently interacting with the ER membrane, tend to accumulate at the perinucleus. Interestingly, such diffusion in a heterogeneous environment follows Chapman’s law rather than the popular Fick’s law. Our findings suggest a novel protein transport mechanism arising solely from characteristics of the intracellular environment.

**Highlights:**

- The interaction of proteins with ER slows down their diffusion at the perinucleus.
- This leads proteins to migrate toward the perinucleus without ATP consumption.
- Frequent ER interaction of larger proteins promotes perinuclear accumulation.
- Diffusion with the physical interaction can be described by Chapman’s law.

**Graphical abstract:** 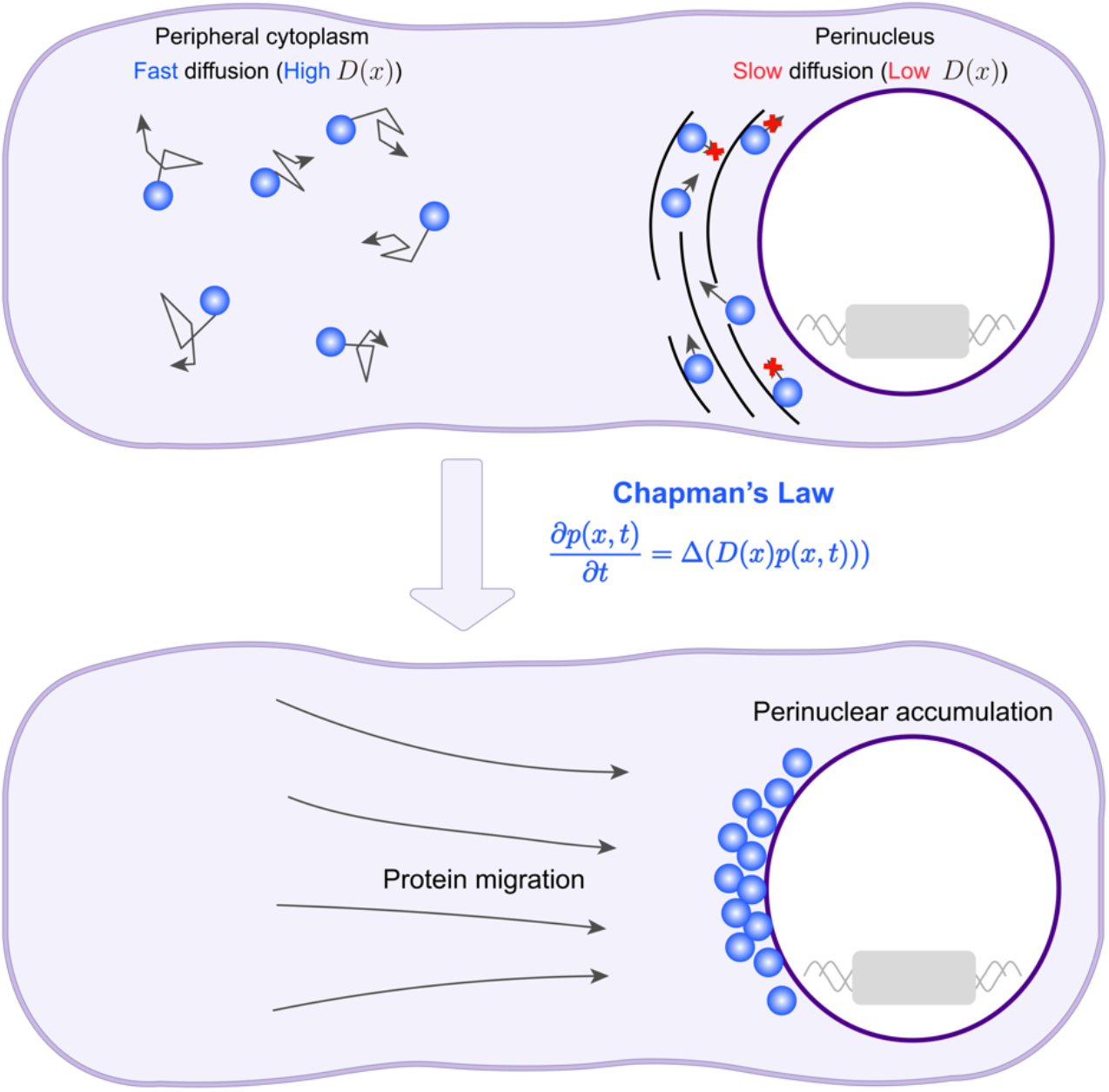

## Introduction

In eukaryotic cells, transcription factors and other proteins that function in the nucleus are translated throughout the cytoplasm but must be translocated across the nuclear membrane^1-3^. The translocation of proteins is regulated by the nuclear pore complexes, which are localized in the nuclear envelope^4,5^. To reach the vicinity of the envelope, nuclear proteins must traverse the cytoplasm. This mainly occurs via intracellular transport mechanisms facilitated by cytoskeletal filament systems, including microtubules and actin, with the assistance of cytoskeletal motors^6-10^. Moreover, actomyosin contraction can also contribute to protein transport by generating cytoplasmic flow^11^. These mechanisms consume adenosine triphosphate (ATP) to fuel motor proteins or contract myosin fibers^6-11^.

Recent studies have shed light on an as-yet-unidentified alternative mechanism that operates passively (i.e., without consuming ATP). Specifically, exogenous particles like nanoparticles and particulate matter, accumulate in the perinuclear region upon entering cells^12-14^. This perinuclear accumulation was also still observed when the energy source (ATP) for the motor proteins was depleted (Figure 1(i))^14^. Moreover, recombinant EGFP-dynactin can migrate toward the nucleus and accumulate in the perinuclear region even without its microtubule binding motif (Figure 1(ii))^15^. Altogether, these studies support the existence of the transport mechanism independent of microtubule-based transport. Interestingly, the accumulation depends on temperature, i.e., decreasing as the temperature decreases (Figure 1(iii))^14^. These characteristics suggest that the unknown mechanism leading to perinuclear accumulation is likely based on diffusion. Notably, diffusion is not homogenous across the cell. In particular, the endoplasmic reticulum (ER), a cellular organelle surrounding the nucleus, is known to impede diffusion^16^. This observation raises the question of whether the heterogeneous diffusive environment in the cytoplasm leads to the migration of proteins toward the nucleus.

**Figure 1.**
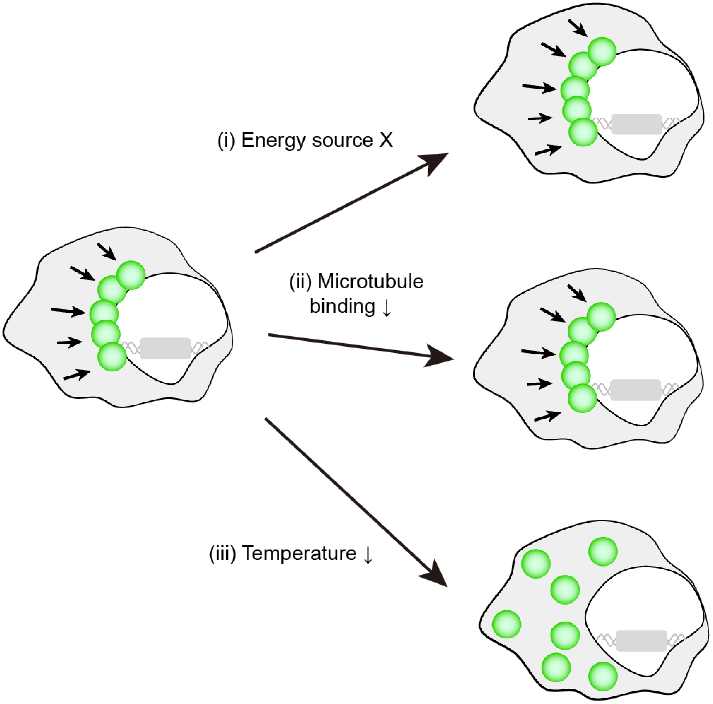
Perinuclear accumulation of protein occurs without consuming ATP. Proteins that need to be transported to the nucleus typically rely on motor proteins powered by ATP for their transport. However, surprisingly, exogenous particle can accumulate in the perinuclear region even when the ATP is depleted (i)^14^, suggesting an alternative transport mechanism. Furthermore, disruption of microtubule binding does not impede this accumulation. Specifically, removing the binding motif of dynactin, a protein assisting dynein in transportation, does not hinder its movement toward the nucleus (ii)^15^. Notably, such accumulation was weakened when the temperature was lowered (iii)^14^. This indicates that the accumulation may have a diffusive character. Reprinted by permission from Rockefeller University Press: Journal of Cell Biology 2007 (doi: 10.1083/jcb.200608128)^15^.

Our study uncovers that the structural characteristics of the intracellular environment that lead to heterogeneous diffusion of proteins also enable proteins to migrate toward the nucleus. Specifically, our agent-based model (ABM) simulation reveals that such migration occurs due to the ER, which impedes protein diffusion. Notably, our ABM also predicts that larger proteins interact with the ER more frequently, leading to their increased accumulation at the perinucleus compared to smaller ones. Indeed, when we analyzed all the proteins whose sizes and subcellular localization are known simultaneously, we found that perinuclear and nuclear proteins tend to have larger radii than cytoplasmic proteins. This passive mechanism offers an energy-efficient alternative for transporting proteins. Furthermore, we found that the diffusion of proteins in a heterogeneous cellular environment can be effectively described using a partial differential equation (PDE) based on Chapman’s law, rather than the conventional Fick’s law that is typically employed to describe diffusion in the cell. This result suggests that employing Fick’s law to explain diffusion in the presence of physical interactions within a cell might not provide accurate results. Taken together, these findings shift our understanding of protein transport, demonstrating that passive mechanisms can be remarkably efficient and that the cell’s own architecture plays a crucial role in guiding protein movement.

## Results

### The cellular environment at the ER leads to perinuclear accumulation

The diffusion process within the cytoplasm is not uniform due to the presence of cellular organelles and macromolecules. Among these organelles, the ER surrounds the nucleus with a maze-like structure (Figure 2A). A previous study suggested that due to the ER, the movement of particles around the nucleus is hindered, and the diffusion significantly slowed down around the nucleus^16^. Furthermore, the diffusion coefficient was decreased as the ER density increased^16^. Intriguingly, when the ER was disassembled, the diffusion coefficient near the nucleus was comparable to that in the peripheral cytoplasm (Figure 2B)^16^.

**Figure 2.**
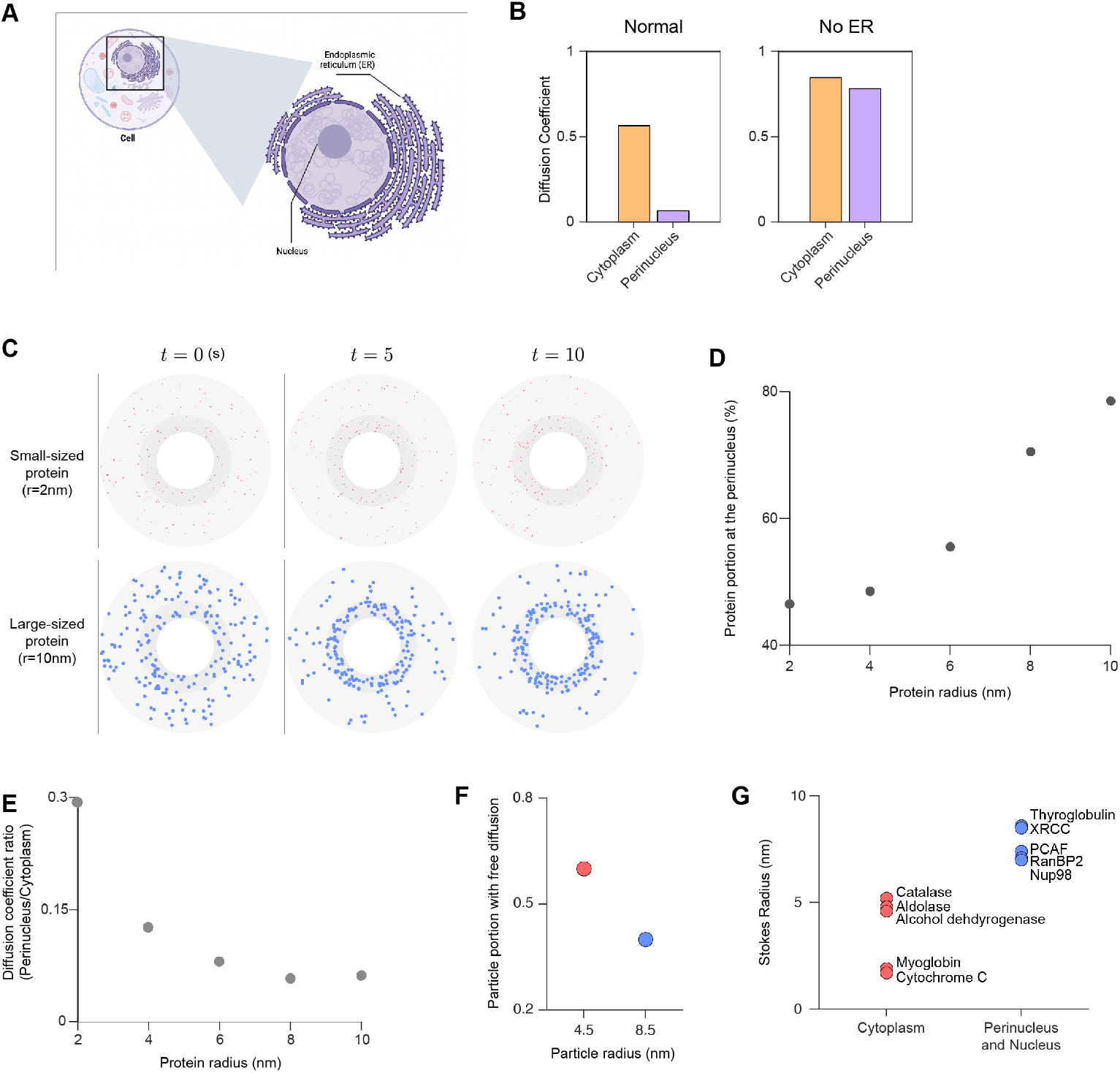
The endoplasmic reticulum (ER) hampers the diffusion of large-sized protein near the nucleus, resulting in migration toward the nucleus. (**A**) The complex structure of ER can impede the diffusion of protein molecules near the nucleus. (**B**) The diffusion coefficient of a quantum dot near the nucleus was much lower compared to the peripheral cytoplasm. When the ER was degraded, the diffusion coefficients became similar throughout the entire cell. The figure is obtained by analyzing Figures 3 and 5 in a previous study^16^. (**C** and **D**) To mimic diffusion in the presence of the ER, an agent-based model (ABM) describing intracellular diffusion was utilized. In particular, we located obstacles with narrow intervals (dark grey region) near the nucleus to describe ER (C). When a protein collided with an obstacle, the protein could bind to the obstacle and become immobile, thereby restricting its movement. This restriction in movement affected larger proteins more than smaller ones since larger proteins more frequently collide with the ER (Figure S1) and thus larger proteins more likely to be accumulate near the nucleus (D). In (C), proteins and obstacles are magnified 20 times for better visualization while biologically realistic sizes are used in the simulation for (D) (see STAR Methods for details). (**E**) The perinuclear diffusion coefficient relative to cytoplasm decreases with increasing protein radius. Protein diffusion coefficients were estimated from their mean-squared displacement (*MSD*) using the Einstein relation (*MSD* = 4*Dt*). (**F**) Consistent with the ABM simulation, diffusion of larger nanoparticles is more likely to be disrupted by the ER compared to that of smaller particles^21^. (**G**) Aligning with the ABM simulation, proteins which are localized in the perinucleus and the nucleus (blue) have larger sizes, while proteins that are localized in the cytoplasm (red) have smaller sizes. The protein size was determined using the Stokes radii (i.e., hydrodynamic radii) from previous studies^27,30-33^.

To explore how proteins move in a heterogeneous diffusive cytoplasm due to the ER, we utilized an ABM, which has previously been used to track molecule movement in various biological systems^17-20^ (see STAR Methods for details). Protein molecules were introduced into the cytoplasm and were set to move randomly in different directions at each time step, simulating diffusive movement. Additionally, we incorporated the ER structure surrounding the nucleus, which impeded protein diffusion. We modeled that this hindrance of protein diffusion stems from protein binding to the ER membrane upon collision with it (see STAR Methods for details).

Interestingly, ABM simulation revealed that as the protein size increases, more proteins accumulate in the perinuclear region (Figure 2C). Specifically, as the radius of protein increases, the portion of protein accumulation at the perinucleus increases (Figure 2D). This is because larger proteins collide more frequently with the ER membrane than smaller proteins despite lower diffusion coefficient of larger proteins (Figure S1). Notably, proteins with a radius of ∼10nm exhibit 2.2 times more collisions in the perinuclear region, compared to proteins with a radius of ∼ 2nm (Figure S1). The higher collision frequency leads to a steeper decline in the diffusion coefficient of larger proteins at the perinucleus compared to their smaller counterparts (Figure 2E), consistent with a previously observed experiment by Reisch et al.: ER impedes the diffusion of nanoparticles more as their size increases (Figure 2F)^21^. Our findings indicate that the ER selectively impedes the diffusion of proteins depending on their sizes, which leads to size-dependent perinuclear accumulation.

To further validate the size-dependent perinuclear accumulation with experimental data, we comprehensively analyzed all the proteins with known sizes and intracellular distribution (Table S1). Specifically, we compared the Stokes radius of proteins^22-34^ based on their subcellular localization, as annotated in previous experiments^35,36^. The proteins localized in the perinucleus or the nucleus have Stokes radii larger than proteins localized in the cytoplasm (Figure 2G), consistent with the model prediction.

### An effective PDE can recapitulate the perinuclear accumulation of proteins

Even though our ABM explains size-dependent accumulation of proteins in perinuclear region (Figure 2), it is challenging to mathematically analyze it. To enable mathematical analysis, we aimed to convert the ABM to a PDE that accounts for the perinuclear accumulation of proteins without explicitly modeling ER geometry. For this, we utilized a simple microscopic model where proteins freely diffuse in the cytoplasm but become immobile upon binding to cellular components, such as the ER (see STAR Methods for details). Under the assumption that the binding and unbinding is faster than the diffusion (see STAR Methods for details), we can derive that this effective diffusion of proteins impeded by the binding can be described by Chapman’s law, i.e., the probability of finding a protein *p*(*x, t*) at a given position *x* and time *t* is governed by the following equation (Figure 3A; see STAR Methods for details):

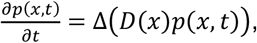

where *D*(*x*) is the effective diffusion coefficient at *x*. Note that Chapman’s law is distinct from the commonly used Fick’s law (Figure 3A).

**Figure 3.**
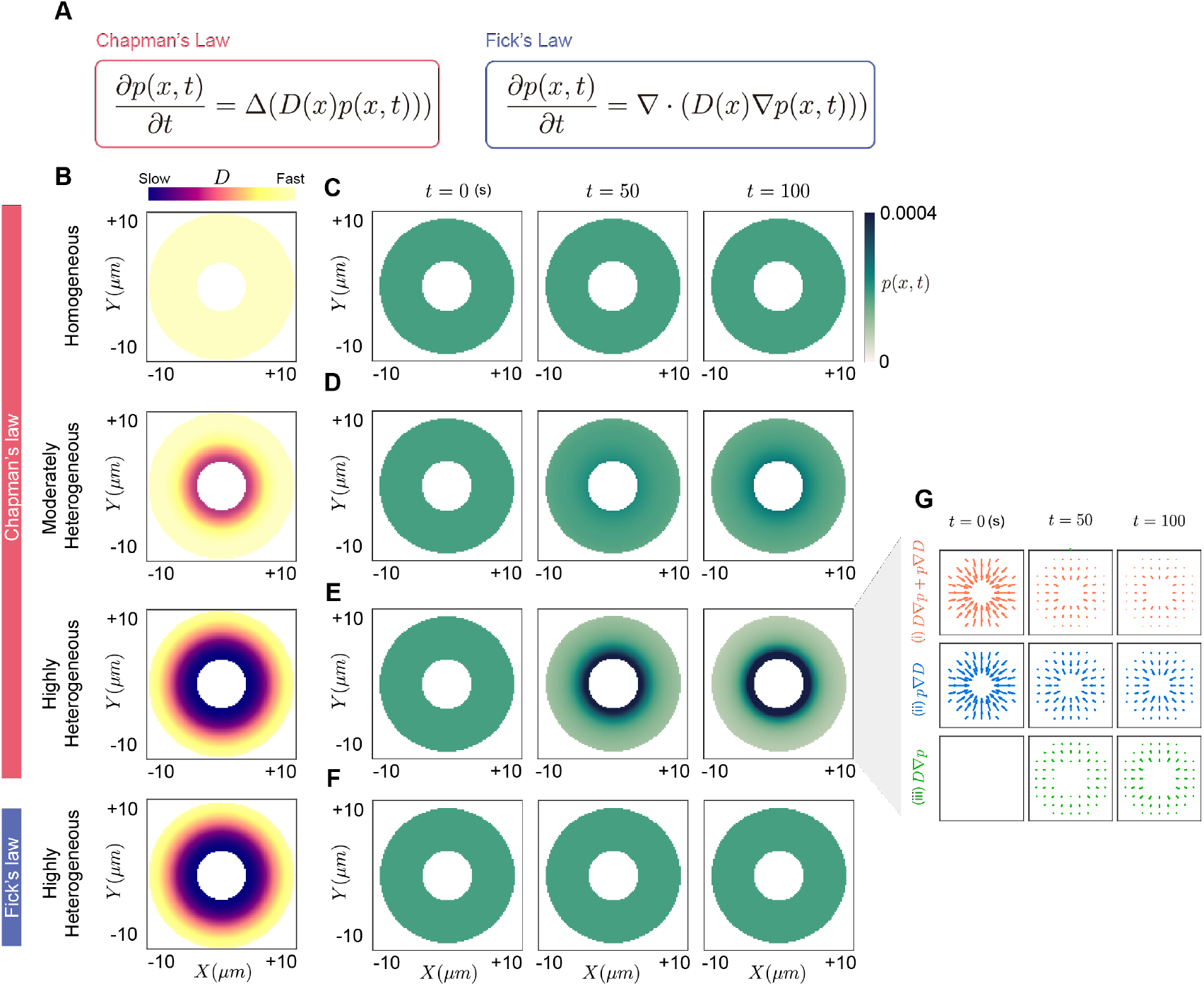
A partial differential equation (PDE) based on Chapman’s law captures the movement of proteins toward the nucleus by heterogeneous diffusion. (**A**) The hindered diffusion due to the physical interaction with the ER near the nucleus can be described by the PDE based on Chapman’s law. This is different from the PDE based on Fick’s law, which has been widely utilized to describe diffusion. Here, *D*(*x*) is the diffusion coefficient at the position *x* and *p*(*x, t*) is the probability of finding the protein at the position *x* and the time *t*. (**B**) Three different environments of diffusion coefficient. (**C-E**) While the PDE based on Chapman’s law resulted in uniform concentration distribution in the homogeneous environment (C), proteins gathered near the nucleus in heterogeneous environments (D and E). (**F**) Fick’s law, which has been widely used to describe diffusion^60-62^, did not exhibit perinuclear accumulation even in the highly heterogeneous environment. (**G**) The flux of Chapman’s law (i) has two components: a drift term (*p*∇*D*) resulting from the gradient of the diffusion coefficient (ii) and a diffusive flux (*D*∇*p*) resulting from the concentration gradient (iii). In the heterogeneous intracellular environment, the drift term (ii) directs proteins toward the nucleus. This leads the diffusive flux (iii) to push proteins outward from the nucleus. While the drift term (ii) remains stronger than the diffusive flux (iii), the protein continues to accumulate near the nucleus (t=0, 50*s*). Eventually, when the drift term (ii) and the diffusive flux (iii) become similar in strength, the net flux reaches the equilibrium, meaning there is no net movement of proteins. (t=100*s*).

The effective diffusion coefficient, *D*(*x*), accounts for how protein diffusion is impeded by physical interactions. In the absence of such interactions, *D*(*x*) remains constant throughout the cytoplasm, resulting in a homogeneous distribution of *D*(*x*) (Figure 3B). Conversely, when protein interact with the ER near the nucleus, *D*(*x*) in this region becomes smaller than that of the peripheral cytoplasm, leading to a heterogeneous distribution of *D*(*x*) across the cytoplasm. Notably, proteins with larger radii more frequently interact with the ER (Figure S1), causing increased hindrance of diffusion compared to proteins with smaller radii (Figure 2E). Thus, *D*(*x*) for larger proteins near the nucleus decreases more than that of smaller proteins. To explore the impact of size-dependent *D*(*x*) on protein movement and accumulation near the nucleus, we investigated moderately and highly heterogeneous *D*(*x*) (Figure 3B).

In the homogeneous environment, perinuclear accumulation did not occur (Figure 3C). On the other hand, in heterogeneous environments, proteins gather near the nucleus (Figures 3D and 3E), resulting in a higher concentration (i.e., higher *p*(*x, t*) near the nucleus). Furthermore, *p*(*x, t*) near the nucleus is higher in the highly heterogeneous environment compared to the moderate heterogeneous environment. Notably, unlike Chapman’s law, the PDE based on conventional Fick’s law cannot capture perinuclear accumulation even in a highly heterogeneous environment (Figure 3F).

To understand why only Chapman’s law, not Fick’s law, leads to perinuclear accumulation, we analyzed the flux (i.e., the amount of protein flows) of PDEs in the highly heterogeneous environment (Figure 3G). In Fick’s law, the flux (*D*∇*p*) is directly proportional to the concentration gradient, causing proteins to migrate from regions of high concentration to regions of low concentration. This flux leads to a steady state with a uniform concentration distribution (Figure S2). In contrast, Chapman’s law incorporates an additional drift term (*p*∇*D*) that arises from the gradient of the effective diffusion coefficient in its flux (*D*∇*p*+ *p*∇*D*) (Figure 3G(i))^37^. Since the drift term heads toward the nucleus, proteins begin to gather near the nucleus from their initial homogeneous concentration distribution (Figure 3G(ii)). As proteins gather near the nucleus, the diffusive flux (*D*∇*p*) directs proteins outward from the nucleus (Figure 3G(iii)), opposing the drift term. Despite this opposition, the drift term (*p*∇*D*) remains stronger than the diffusive flux (*D*∇*p*), causing the total flux (*D*∇*p* + *p*∇*D*) to still be directed toward the nucleus (Figure 3G(i)). When the diffusive flux cancels out the drift term, the total flux becomes zero, resulting in a steady state with protein accumulation near the nucleus (Figure 3G(i)). At the steady state (i.e., *D*∇*p* + *p*∇*D* = ∇(*Dp*) = 0), *p*(*x, t*) is inversely proportional to *D*(*x*), highlighting the crucial role of physical interactions in perinuclear accumulation. Taken together, our findings suggest that Chapman’s law is more suitable than Fick’s law for describing the existing experimental results on diffusion in cellular environments^36^.

### An ABM, which corresponds to PDE, accurately captures the perinuclear accumulation

Although the PDE based on Chapman’s law provides a simple way to account for the perinuclear accumulation, it can become cumbersome for complex cell shapes and intracellular environments. In such cases, using ABM equivalent to PDE based on Chapman’s law can be a versatile alternative. Specifically, incorporating heterogenous effective diffusion in ABM without explicitly modeling binding and unbinding reactions unlike the ABM in Figure 2 can be an efficient way to simulate perinuclear accumulation of proteins within a heterogeneous diffusive environment. Thus, we aimed to explore the conditions under which ABM becomes equivalent to the PDE based on Chapman’s law, enabling the reproduction of perinuclear accumulation of proteins within a heterogeneous diffusive environment (Figure 3A).

In ABMs, the diffusion of proteins is characterized by a random walk^18,20^. The step size of each step of random walk is proportional to the square root of the effective diffusion coefficient^38^. Since the effective diffusion coefficient varies within the cytoplasm (Figure 3B), the effective diffusion coefficient can change even during a single step. Thus, different choices of step size can be made based on the position where the effective diffusion coefficient is determined. For example, determination of the effective diffusion coefficient can be based on the current position before the step or the position after the step. The choice of where to evaluate the diffusion coefficient results in distinct types of random walk, leading to distinct dynamics. Notably, when the effective diffusion coefficient is evaluated at the current position, the random walk becomes equivalent to the discretization of the Ito integral, and the dynamics of ABM follows Chapman’s law^39^ (Figure 4A). Therefore, when this random walk scheme was incorporated into the ABM, perinuclear accumulation was observed over time in a highly heterogeneous diffusion environment (Figure 4B). Taken together, ABM utilizing the effective diffusion coefficient of current position of the protein can effectively simulate protein diffusion in heterogeneous intracellular environments to capture perinuclear accumulation of proteins.

**Figure 4.**
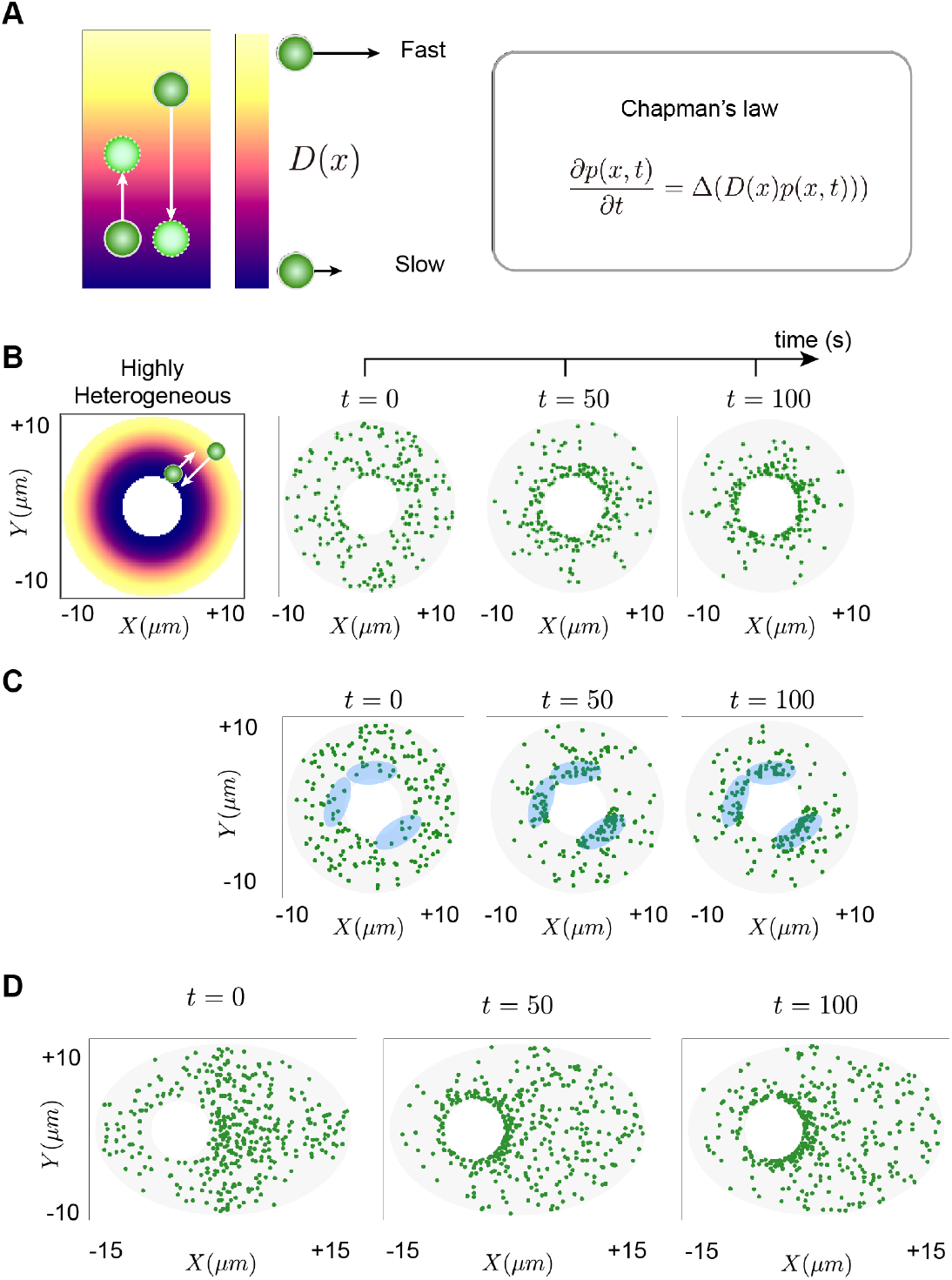
ABM scheme that corresponds to Chapman’s law results in the perinuclear accumulation of proteins. (**A**) For the ABM simulation, the step size of the protein movement is determined by the diffusion coefficient. When the diffusion coefficient varies over the cytoplasm, the step size can vary depending on the position at which the diffusion coefficient is obtained. When the step size is determined by the molecule’s current position, the dynamics of proteins is equivalent to Chapman’s law. (**B**) This ABM scheme was simulated in the highly heterogeneous environment, which is used in Figure 3E. Initially (t=0*s*), proteins were uniformly dispersed throughout the cytoplasm. Over time, the proteins gradually accumulated near the nucleus (t=50, 100*s*). (**C**) To simulate the ER partially surrounding the nucleus, we selectively reduced the diffusion coefficient in specific perinuclear regions (blue). Even when the ER is partially surrounding the nucleus, perinuclear accumulation still occurred. (**D**) Besides, ABM provided a convenient approach for describing diffusion in cells of various shapes, including elliptical cells. Proteins again accumulated near the nucleus in elliptical cells.

Such ABM framework can depict the diffusion in various intracellular environments. Specifically, the ABM can be easily adapted to describe more realistic but complex environments, such as cells with the ER structure partially surrounding the nucleus (Figure 4C), or elliptical-shaped cells (Figure 4D) (see STAR Methods for details). Remarkably, even in these complex cell geometries, the ABM simulations replicate perinuclear accumulation (Figures 4C and 4D) observed in idealized cells (Figure 4B).

## Discussion

To maintain normal physiological processes within cells, it is crucial to effectively and selectively transport to the nucleus proteins translated in the cytoplasm. Our research has shown that proteins can passively diffuse to accumulate near the nucleus without requiring active transport mechanisms, such as transport by motor proteins along microtubules. This accumulation occurs because physical interactions between proteins and the ER slow the diffusion at the perinucleus relative to the peripheral cytoplasm (Figures 2E and 3B). Larger proteins are more likely to accumulate at the perinucleus (Figures 2C and 2D) because physical interactions occur more frequently as the size of the molecules increases (Figure S1). Our findings reveal that diffusion within a heterogeneous cellular environment enables the ATP-free transport of large-sized proteins, contrasting with the conventional microtubule-based transport, which requires ATP as an energy source^6,7^.

In addition to the ER, the perinucleus is more crowded than the cell periphery because the perinucleus contains more organelles and denser microtubule networks^40^. This molecular crowding can also impede protein diffusion if proteins can bind to organelles or microtubules. Since physical interactions occur more frequently as the size of the molecules increases (Figure S1), larger proteins are more likely to accumulate at the perinucleus (Figures 2C and 2D). This provides insight for previous experimental observations: 1) proteins destined to reach the nucleus often form complexes before translocating to the nucleus^41,42^, 2) attaching specific moieties to anticancer drug particles, thereby increasing their size, facilitates targeting to the ER^43,44^, and 3) different types of particles, such as lipid nanoparticles and CaCO3, tend to accumulate in the perinuclear region based on their sizes^12-14^. While we focused on proteins with sizes comparable to the ER width (∼20 nm), cells also contain significantly larger particles (>>20nm) like vesicles and lysosomes^45,46^. These extremely large particles might interact with actin filaments, whose mesh size is approximately 50nm^40^. Investigating the diffusion of these extremely large particles and their potential interactions with the cytoskeleton would be an interesting avenue for future work.

The ER consists of two main structural components: a tubular network structure in the cell periphery and a sheet structure near the nucleus. While both structures contribute to the heterogeneity in diffusion coefficients of proteins^16^, their effects differ significantly. The tubular network results in the compartmentalization of the diffusion in the cell periphery, and the diffusion coefficient changes minimally due to a sparse ER density^16^. In contrast, the sheet structure is highly dense near the nucleus, and thus large proteins slow down as they approach the nucleus (Figure 2E)^16^. This suggests that the sheet structure is important for helping proteins move towards the nucleus, while the tubular network likely has little effect on translocation.

Among various types of PDEs describing diffusion^47-55^, it is challenging to determine a priori the most appropriate diffusion law (e.g., Fick’s law or Chapman’s law) for modeling diffusion in heterogeneous environments^56,57^. This challenge arises from the diverse physical causes giving rise to heterogeneous diffusion, each manifested in distinct forms of diffusion equations^57^. Consequently, a thorough understanding of the underlying physical phenomena governing heterogeneity is essential. Nevertheless, intracellular diffusion has been primarily described by Fick’s law^58-62^. Our study suggests that describing protein diffusion with binding to cellular components like the ER (Figure 3B) is equivalent to the “trap model” in Sokolov, resulting in Chapman’s law^57^. When such cellular components impede the diffusion near the nucleus, the PDE based on Chapman’s law but not Fick’s Law yields a non-uniform concentration distribution in a steady state, and accurately describes the perinuclear accumulation (Figures 3C-3F). However, mathematical models describing circadian clocks have adopted Fick’s law to describe the diffusion of clock proteins^18,59^, although a recent study showed that a clock protein is accumulated at the perinucleus^18^. Thus, utilizing Chapman’s law to develop the circadian clock model would be more appropriate to investigate the spatiotemporal dynamics of molecular circadian clock^18,20,59,63-65^. Furthermore, it would be an interesting future work to investigate dynamics of various biological oscillators based on transcriptional feedback loops requiring nuclear transport of transcriptional factors (e.g., p53 oscillator^66-68^, NF-*k*b oscillator^69^, and synthetic oscillator^70-73^) when Chapman’s law is used rather than Fick’s law.

Previous studies of diffusion in heterogeneous environments, particularly focusing on highly crowded cellular environments, have highlighted the prevalence of subdiffusion^74-76^ for various particles, including small labeled proteins, lipid and insulin granules, and virus particle^77-80^. This subdiffusive behavior, characterized by slower-than-linear growth in displacement variance, suggests hindered transport and potential inefficiencies in cellular processes. On the other hand, our study demonstrates how heterogeneous diffusion, specifically under the influence of physical interactions with the ER, can facilitate directional protein transport toward the nucleus, as implied by Chapman’s law. Consistently, heterogeneous diffusion was known to be beneficial for reducing the first-passage time of reaction events, even though it slows down the overall dynamics of proteins^81^. Taken together, diffusion in heterogeneous environments can also contribute to efficient cellular functioning.

### Limitations of the study

We have comprehensively examined all the proteins for which both their sizes and subcellular localizations (i.e., locations of proteins) are known, and this analysis supports that larger proteins tend to accumulate near the nucleus (Figure 2G). However, the scarcity of proteins with known sizes and subcellular localizations restricts the generalizability of our conclusions. It would be an important topic to explore how subcellular localization and proteins sizes are correlated across a broader range of proteins. Moreover, it is crucial to verify whether a protein does not contain NLS when examining its subcellular localization, since proteins and particles with NLS are reported to induce microtubule-based transport^82,83^. To further strengthen our conclusion, it would also be helpful to investigate whether smaller proteins can accumulate near the nucleus after fusing with fluorescent proteins, as these modifications can significantly increase the size of protein^84^.

## Acknowledgements

We thank Yong Jung Kim and Ho-Youn Kim for the helpful discussion. We thank Life Science Editors for editorial assistance. This work was supported by the Institute for Basic Science (grant no. IBS-R029-C3) (J.K.K.) and Welch Foundation grant no. C-1995 (O.A.I.). Some parts of figure 2 were retrieved from Biorender.

## Author Contributions

S.J.C., D.W.K., S.L. and J.K.K. designed the study. S.J.C. performed computational modeling and simulation and all authors contributed to analysis. J.K.K. supervised the project. S.J.C. and J.K.K. wrote the manuscript draft, and all authors revised the manuscript.

## Declaration of interests

The authors declare no competing interests.

## STAR★Methods

### RESOURCE AVAILABILITY

#### Lead contact

Further information and requests for resources and reagents should be directed to and will be fulfilled by the lead contact, Jae Kyoung Kim.

#### Materials availability

This study did not generate new unique reagents.

#### Data and code availability

- This paper analyzes publicly available existing data. These accession numbers for the datasets are listed in the key resources table.
- The MATLAB and Netlogo codes of the computational package are available in the following Database: The link will be available upon acceptance of the manuscript.
- Any additional information required to analyze the data is available from the lead contact upon request.

### METHOD DETAILS

#### ABM describing diffusion in the presence of the ER

To describe the heterogeneous diffusion in a cell, we utilized an ABM which describes the intracellular diffusion of proteins. Following previous studies^18,20^, we simplified the cell geometry as a two-dimensional circular shape with the radius of 10µm^85^. Within the cell, a circular nucleus resided, sharing the same center and possessing a radius that is one-third of the cell radius (i.e., 3.33µm), so that the nucleus occupies about 10% of the cell area^18^. Furthermore, we incorporated the ER by introducing obstacles near the nucleus (Figure 2C). These obstacles were distributed across 80 concentric layers. Starting from 3.58µm, each layer increased in radius by 0.25µm, reflecting the width of the ER sheet structure^85^. Within each layer, 1,000 uniformly distributed obstacles were uniformly distributed to mimic increased ER density as proteins approached the nucleus. Each obstacle was modeled as a circle with a 0.0025µm radius.

A protein molecule was represented as a circle with the radius *r*. In our simulation, we varied *r* between 2 nm, corresponding to the radius of carbonic anhydrase^33^, to 10 nm, corresponding to the PER2 complex^86^. The diffusion of proteins was described by using a random walk. During each time step of length *Δt*, each molecule was moved in a randomly chosen direction with a step size *δ*. The angles of movement were randomly sampled from the uniform distribution, ranging from 0° to 360°. The step size *δ* was chosen to be sufficiently smaller than the protein size to ensure precise and fine-scale movements. For example, for a protein with 10nm radius, *δ* was set to be 1.6nm. For proteins with smaller radii, the step size was adjusted by using the Stokes-Einstein equation (i.e.,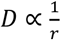 and thus 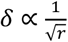). The time step length (*Δt*) was determined by using the Einstein relation *δ*^2^ = *4DΔt*^*38*^, where *D* is the diffusion coefficient. In this study, we utilized the known diffusion coefficient of PER2 protein (*D=*0.2µ*m*^2^*/s*)^87^ and its radius of 10nm^86^, (i.e., *δ* = 1.6nm), resulting in *Δt* = 0.0000032*s*.

When proteins collided to an obstacle, they were repositioned to the point of collision. Subsequently, they bound to the obstacle with a binding probability of 1 − exp(−*k*_*bind*_*Δt*), where *k*_*bind*_ is the binding rate. Once bound, at each time step, proteins could unbind with a probability of 1 − exp(−*k*_*unbind*_ *Δt*), where *k*_*unbind*_ is the unbinding rate. In our simulation, *k*_*bind*_ and *k*_*unbind*_ were set to 1,562,500*/s* and 625*/s*, respectively.

We implemented reflecting boundary conditions to prevent the protein from leaving the cytoplasm. That is, we repositioned proteins as if they were elastically reflected by the boundary (cell membrane or the nuclear membrane). Specifically, proteins exceeding the cytoplasmic boundary (cytoplasmic membrane or nuclear membrane) are mirrored across the tangent line of the boundary at the collision point with the boundary. Additionally, for the initial condition, protein began diffusion from a uniform distribution of the protein throughout the cytoplasm.

#### Physical interpretation of Chapman’s law

Since our mechanism of protein transport operates without the need for energy sources, such as ATP hydrolysis, we expect that steady state distribution will correspond to the thermodynamic equilibrium. This steady state with non-uniform concentration distribution should result from the energy difference between spatial locations. We postulate that the difference in energy level comes from the physical interaction between proteins and the cellular organelles, including the ER.

To investigate this, we considered a one-dimensional diffusion of protein in the cytoplasm. Specifically, we considered a protein moving along a linear chain of *N* sites (*i* = 1*∼N*), where the chain represents the cytoplasm and each site is separated by an interval *h*. The protein located at the site *i* can interact with cellular organelles, such as the ER, with a binding rate of 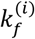 and it can also detach from the organelles with an unbinding rate of 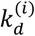. If the protein at the site *i* is not bound, it can diffuse to the adjacent site (e.g., *i* + 1 or *i* − 1) with a diffusion coefficient of *D*. Denoting the probability of finding the bounded and unbounded protein at the site *i* at time *t* by 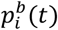 and 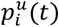, respectively, we obtain the following ordinary differential equation (ODE):

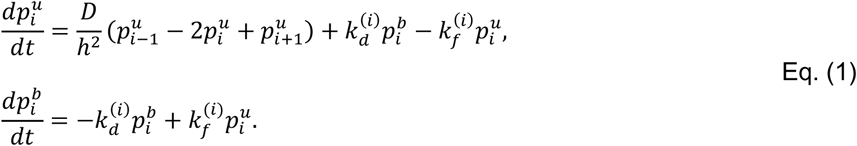

Here, we use the convention of 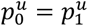 and 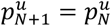 by imposing Neumann boundary conditions. Since we are primarily interested in the probability of finding proteins at the i-th site, regardless of whether they are bound or not, we introduce a new variable 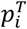 the probability of finding the protein at the site i, i.e., 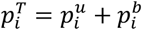. Then, Eq. (1) can be written as shown below:

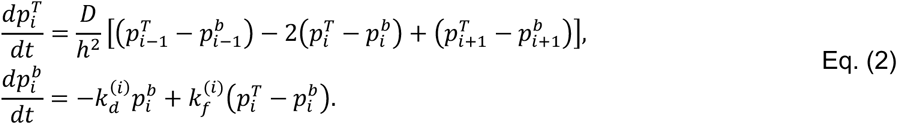

We assume that binding and unbinding reactions occur significantly faster than diffusion. Then 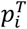 is affected by only slow diffusion, whereas 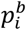 is affected by fast binding and unbinding reactions, leading to a complete time scale separation. This allows us to assume that 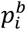 reaches the quasi-equilibrium (the fast binding and unbinding equilibrium) at each site before 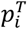 changes appreciably (i.e., 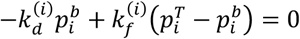 for each *i*)^88-92^. Then, 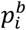 can be expressed by 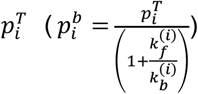 Consequently,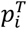 follows a reduced ODE shown below:

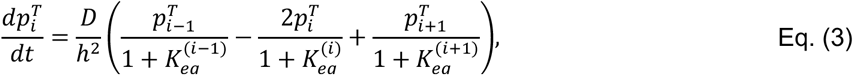

where 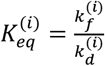 Taking the limit of infinitesimal interval (*h* → *0*), we obtain the following Chapman’s law:

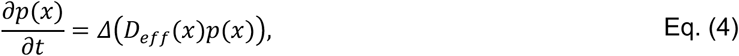

where 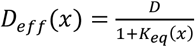 To validate the model reduction, we compared whether the original model (Eq. (1)) and the reduced model exhibit similar behavior. As long as the binding and unbinding occur significantly faster than the diffusion, the two models exhibit similar dynamics (Figure S3).

#### Numerical simulation for the PDE

After deriving the PDE based on Chapman’s law in the previous section, we solved the PDE numerically. Depicting diffusion within a cell accurately using the standard finite difference method on a rectangular domain presents challenges because cells usually have a non-rectangular shape and contain a nucleus where diffusion does not occur. To address this issue, we adopted an approach from a previous study which described the diffusion under a domain with a complex geometry^93^. Specifically, we solved the PDE based on Chapman’s law 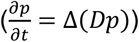 in a rectangular domain Ω = [−*L, L*] × [−*L, L*], where, *L* was set to be 1.1 ⋅ *R*_*c*_, where *R*_*c*_ is the radius of the cell. By letting 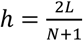 we obtained a uniform mesh with a set of grid edges as Ω^*h*^ ={(*x*_*i*_, *y*_*j*_): 1 ≤ *i, j* ≤ *N*}, where *x*_*i*_ = −*L* + (*i* – 1)*h* and *y*_*j*_ = −*L* + (*j* − 1)*h*. Here, we used *N* = 100. At the edge (*x*_*i*_, *y*_*j*_), the boundary control function *G*_*-i,j*_ was defined as 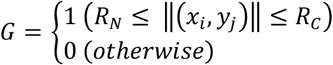 where *R*_*N*_ is the radius of the nucleus. In the grid edge, *G* is defined as 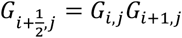 and 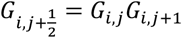 Utilizing *G* allowed us to utilize the finite difference method in a complex domain mimicking the cell by simply computing PDE in a rectangular domain^93^.

We denote the discrete approximation *p*(*x*_*i*_, *y*_*j*_, *nΔt*) as 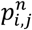. Then, we can define the discrete differential operator ∇_*d*_ as follows:

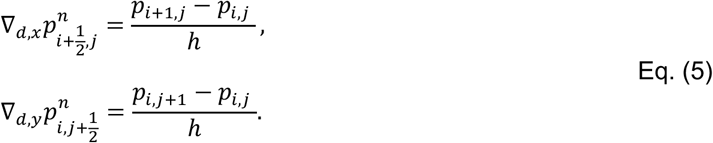

Furthermore, to calculate ∇_*d*_ at the boundary, the value of 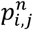 at ghost points located outside of the domain also required (*i, j* = 0 *or N* + 1). To determine values out of the domain, we utilized the Neumann boundary condition. Specifically, we set 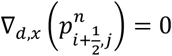 for *i* = 0, *N* + 1 and 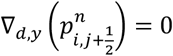 for *j* = 0, *N* + 1. The boundary condition and boundary control function *G* ensures the conservation of *p*(*x, t*) over time, i.e., ∫ *p*(*x, t*)*dx* remains constant over time^94^. Finally, the PDE based on Chapman’s law can be written as the following:

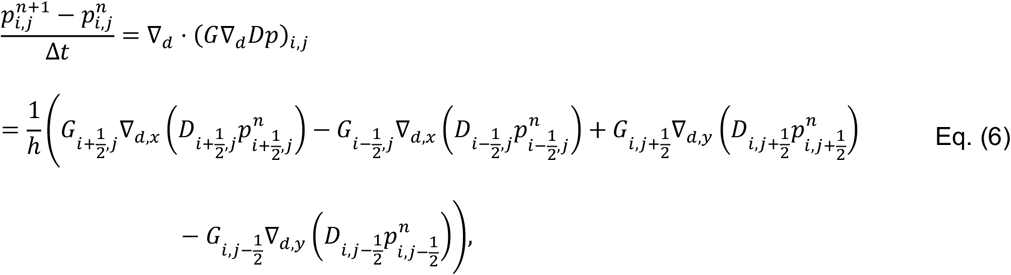

for 1 ≤ *i, j* ≤ *N*, where *h*=0.2µm and *Δt* = 0.00004s were used. To determine *G*, the same value of *R*_*N*_ and *R*_*C*_ as in the ABM are used. Using the PDE (Eq. (6)), we obtained the solution of Chapman’s law in Figure 3.

To describe the diffusion with the existence of physical interactions, we have utilized various effective diffusion coefficients at (*x, y*), *D*(*x, y*). A homogeneous case described by using *D*(*x*) = *0*.2µm^2^*/s*. For moderately and highly heterogeneous cases, we employed functions that decrease as the protein approaches the nucleus. Specifically, we used *D*(*x, y*) = 0.2 · 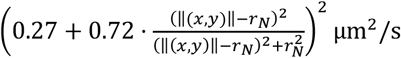 and 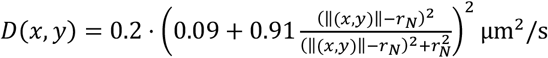 to describe moderately heterogeneous and highly heterogeneous cases, respectively. Here, *r*_*N*_ is the radius of the nucleus, and ‖(*x, y*)‖ is the distance between the protein and the cell center.

#### Simulating ABM based on Chapman’s law

To simulate an ABM equivalent to Chapman’s law, we introduced a variation in the protein’s movement (Figures 4B-4D). Specifically, the protein molecule at position (*x, y*) moves by a step size *δ*(*x, y*) in a random direction. The step size of the protein was determined by the diffusion coefficient of the current position. We modeled the decrease in the step size by using a Hill function: 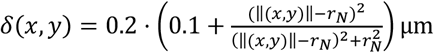µm, where *r*_*N*_ is the radius of the nucleus, and ‖(*x, y*)‖ is the distance between the protein and the cell center. This choice of step size of the same form of Hill function to PDE (*δ*^*2*^ *α D*) ensures that the ABM aligns with the highly heterogeneous environment. As the protein approaches the nucleus, the step size decreases to be 10% of the maximum value (=0.02µm). The length of time of step Δ*t* was obtained by using the relation 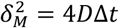, where *δ*_*M*_ is the maximum value of the step size and *D* is the diffusion coefficient of PER2 protein (*D=*0.2µm^*2*^*/s*)^87^.

In Figures 4C and 4D, we have extended the ABM to describe cells with diverse geometries. To model the ER structure partially surrounding the nucleus in Figure 4C, we implemented the following step size function: 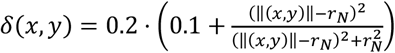. 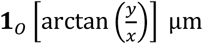 where *O* = {*θ*: 6*0*° ≤ θ< 21*0*° or 27*0*° ≤ θ< 36*0*°} is and 1_*0*_(*x)* is the indicator function of *O*. In Figure 4D, a cell was represented by an ellipse with a semi-major axis of 15µm and a semi-minor axis of 10µm, centered at the origin (Figure 4D). The nucleus was depicted as a circle with a radius of 3.33µm, centered at (−5µm, 0µm).

## Notes

### Competing Interest Statement

The authors have declared no competing interest.

### Summary of Updates

Error in figure is revised

